# Ancestry-agnostic estimation of DNA sample contamination from sequence reads

**DOI:** 10.1101/466268

**Authors:** Fan Zhang, Matthew Flickinger, InPSYght Psychiatric Genetics Consortium

**Affiliations:** Center for Statistical Genetics, University of Michigan, Ann Arbor, MI; Department of Computational Medicine and Bioinformatics, University of Michigan Medical School, Ann Arbor, MI; Department of Biostatistics, School of Public Health, University of Michigan, Ann Arbor, MI

## Abstract

Detecting and estimating DNA sample contamination are important steps to ensure high quality genotype calls and reliable downstream analysis. Existing methods rely on population allele frequency information for accurate estimation of contamination rates. Correctly specifying population allele frequencies for each individual in early stage of sequence analysis is impractical or even impossible for large-scale sequencing centers that simultaneously process samples from multiple studies across diverse populations. On the other hand, incorrectly specified allele frequencies may result in substantial bias in estimated contamination rates. For example, we observed that existing methods often fail to identify 10% contaminated samples at a typical 3% contamination exclusion threshold when genetic ancestry is misspecified. Such an incomplete screening of contaminated samples substantially inflates the estimated rate of genotyping errors even in deeply sequenced genomes and exomes.

We propose a robust statistical method that accurately estimates DNA contamination and is agnostic to genetic ancestry of the intended or contaminating sample. Our method integrates the estimation of genetic ancestry and DNA contamination in a unified likelihood framework by leveraging individual-specific allele-frequencies projected from reference genotypes onto principal component coordinates. We demonstrate this method robustly and accurately estimates contamination rates across different populations and contamination rates. We further demonstrate that in the presence contamination, quantitative estimates of genetic ancestry (e.g. principal component coordinates) can be substantially biased if contamination is ignored, and that our proposed method corrects for this bias. Our method is publicly available at http://github.com/Griffan/verifyBamID

## Introduction

Sample contamination is a common problem in DNA sequencing studies. Contamination may occur during sample shipment (due to spillage across wells, pipetting errors, insufficient dry ice), library preparation (due to gel cut-through in fragment size selection or unexpected switch between barcoded adaptors *in*-*vitro*), *in*-*silico* demultiplexing from a sequenced lane into barcoded samples, or on many other unexpected occasions. Even modest levels of contamination (e.g. 2-5%) within a species substantially increase genotyping error, even for deeply sequenced genomes^1^. Accurate estimation of DNA contamination rates allow us to identify and exclude contaminated samples from downstream analysis, and genotypes of moderately contaminated samples (e.g. <10%) can be improved by accounting for contamination in genotype calling^1^.

Previously we developed methods and a software tool, *verifyBamID*^2^, to estimate DNA contamination from sequence reads given known population allele frequencies of common variants. Many investigators and most major sequencing centers use *verifyBamID* as a part of their standard sequence processing pipeline. However, we have shown that *verifyBamID* can underestimate DNA contamination rates if the assumed population allele frequencies are inaccurate^2^. Such an underestimation can be avoided if correct population allele frequencies are provided in an ideal circumstances. However, in early stage of sequence analysis, performing a tailored customization of quality control (QC) steps for each sequenced genome based on their ancestry is not always feasible or or sometimes impossible. Such a tailored customization requires planned coordination between sequencing centers and study investigators prior to sequencing to share the self-reported ancestry (which is not always accurate) or estimated ancestry from external genotypes (which is not always available). Modifying the QC pipeline to accommodate study-specific or sample-specific parameters may not be a possible option for large sequencing centers. Even if such a tailored customization of QC pipeline is possible, preparing per-sample ancestry prior to QC may delay time-sensitive issues in the sequencing procedure. If contamination rates can be accurately estimated without having to know the ancestry or allele frequencies a priori this will simplify the sequence analysis pipeline and expedite the QC.

Here we describe a novel method to robustly detect and estimate DNA contamination by modelling the probability of observed sequence reads as a function of “individual-specific allele frequencies” that account for genetic ancestry of a sample. Instead of assuming that the population allele frequencies are known, we represent individual-specific allele frequencies as a function of genetic ancestry using principal component coordinates and the reference genotypes from a diverse population, e.g. Human Genome Diversity Project (HGDP)^3^ or 1000 Genomes^4^. We then jointly estimate genetic ancestry and contamination rates of a sequenced individual based on a mixture model, without requiring the assumption that population allele frequencies are known.

Our method enables accurate ancestry-agnostic estimation of contamination through a unified likelihood framework that incorporates genetic ancestry and contamination together. We show that our method provides (1) comparable or more accurate estimates of genetic ancestry than existing methods such as *TRACE/LASER^5,6^* even in the absence of contamination and (2) reduced bias in contamination rate estimates compared to our previous method requiring known population allele frequencies using *in silico* contaminated datasets and sequenced genomes from the InPSYght psychiatric genetics sequencing study.

## Material and Methods

We aim to jointly estimate sample contamination rates and genetic ancestry from sequence reads without specifying population allele frequencies. First, we describe our previous mixture model to estimate contamination rates assuming population allele frequencies are known. Second, we introduce a model for sequence reads using population allele frequencies as a function of genetic ancestry represented in principal component coordinates. Third, we extend the model to enable joint estimation of contamination rates and genetic ancestry. Fourth, we evaluate our methods using *in silico* contaminated samples and whole genome sequence data from the InPSYght study.

### Likelihood-based mixture model for DNA sequence contamination

In our previous contamination detection methods^2^, we assumed that the DNA sequence reads from an intended sample are contaminated by sequence reads from at most one contaminating sample from the same population, and that the population allele frequencies of all analyzed genetic variants are known. For each bi-allelic variant *i* (1 ≤ *i* ≤ *m*), let *b_ij_* ∈ {*R*, *A*, *O*} (1 ≤ *j* ≥ *D_i_*) be the observed base call representing the reference allele (R), alternate allele (A), or other allele (O) for the *j*-th read that overlaps the variant; *D_i_* is the observed sequence depth at variant *i*. Let *e_ij_* ∈ {0,1} be a random variable indicating whether a sequencing error did (1) or did not (0) occur for observed base *b_ij_*; we assume *e_ij_* follows a Bernoulli distribution with success probability
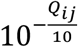 where *Q_ij_* is a phred-scale base quality score of *b_ij_*. In the absence of contamination, if the true genotype *g_i_* ∈ {0,1,2} represents the count of alternate alleles of the sequenced sample, then
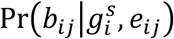 can be easily represented as in Table 1, making the simplifying assumption of equally likely errors across four possible nucleotides.

**Table 1.** Conditional probability P(b_ij_ | g_i_, e_ij_) of read b_ij_ given true genotype g_i_ and the variable representing the event of base calling error e_ij_, as described in (Jun et al 2012^2^)

| True Genotype $g_i$ | Base Calling Error Event $e_{ij}$ | $\Pr(b_{ij} = R)$ | $\Pr(b_{ij} = A)$ | $\Pr(b_{ij} = O)^b$ |
| --- | --- | --- | --- | --- |
| $g_i = RR^a$ | $e_{ij} = 0$ | 1 | 0 | 0 |
| | $e_{ij} = 1$ | 0 | 1/3 | 2/3 |
| $g_i = RA^a$ | $e_{ij} = 0$ | 1/2 | 1/2 | 0 |
| | $e_{ij} = 1$ | 1/6 | 1/6 | 2/3 |
| $g_i = AA^a$ | $e_{ij} = 0$ | 0 | 1 | 0 |
| | $e_{ij} = 1$ | 1/3 | 0 | 2/3 |

<sup>a</sup> RR, RA, AA: homozygous reference, heterozygous, and homozygous non-reference genotypes
<sup>b</sup> O: alleles other than R or A; assumes four possible alleles (bases)

We assume that the observed sequence reads are a (1 – α): α mixture of intended and contaminating reads given a contamination rate 0 ≤ α ≤ 1. Let
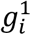 and
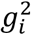 represent the true genotypes of the intended and contaminating samples at variant *i*, respectively. Then the mixture model likelihood of each observed base becomes

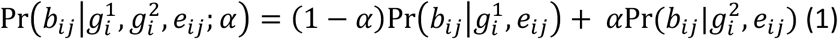

Assuming a homogenous population with known population allele frequency *f_i_* and Hardy-Weinberg Equilibrium (HWE),
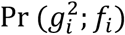 follows a Binomial(2, *f_i_*) distribution. Under the simplifying assumption of independent variants, the likelihood of the contamination rate becomes

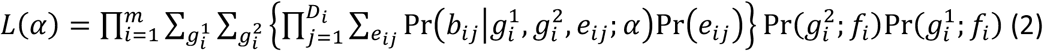

The maximum likelihood estimate (MLE) of contamination rate
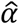 can be obtained using Brent’s algorithm^7^.

As we previously reported^2^, this model assumes correctly specified population allele frequencies *f_i_*.

### Likelihood-based estimation of genetic ancestry (in the absence of contamination)

We extend this model to incorporate genetic ancestry. The key idea of this extension is to use the individual-specific allele frequency (ISAF)^8,9^ to model the likelihood of the sequence reads. Several methods, including Spatial Ancestry Analysis (SPA)^10^ and logistic factor analysis (LFA)^9^, previously proposed modelling allele frequency as a function of genetic ancestry via principal component (PC) coordinates.

Let *G* be an *m* × *n* genotype matrix (where *g_ij_* = 0, 1, or 2 is the number of non-reference alleles at variant *i* in individual *j*) of a genetically diverse reference panel of size n, such as 1000 Genomes or HGDP. We define ISAF *f_i_* (0 ≤ *f_i_* ≤ 1) for variant *i* as a weighted average of genotypes from the reference panel
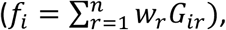 where 0 ≤ *w_r_* ≤ 1 and *G_ir_* ∈ {0,1,2} for individual *r*. For a homogenous population,
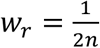 results in a *pooled allele frequency* across all individuals in the reference panel. If each individual can be categorically represented as a one of *k* mutually exclusive subpopulations, the *population*-*specific allele frequency* for the subpopulation *s* ∈ {1,2, ⋯, *k*} can be represented as
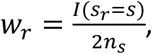 where and *s_r_* ∈ {1,2, ⋯,*k*} represents the subpopulation that individual *<u>r</u>* belongs to, and *n_s_* represents the size of subpopulation *s*. More generally, if individual’s genetic ancestry is represented as continuous variables (such as PCs, SPAs, or LFAs), the individual-specific allele frequency (ISAF) can be represented as a function of the continuously represented genetic ancestry^9,5^.

The estimated ISAF can be viewed as (one half times the) genotype dosages approximated from a fixed number(=*K*) of factors, such as PCs, SPAs, or LFAs. In our method, we used a linear model to estimate ISAF from PCs, similar to previous studies^8,9^. Given the reference panel genotype matrix *G*, let
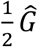 be the *ISAF matrix* as a function of top *K* factors. ISAF matrix
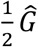 should well approximate
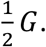 For example, under a linear model typing principal component analyst takes the singular value decomposition (SVD) of the mean-centered genotype matrix
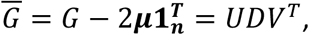 where
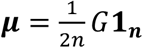 is the pooled allele frequencies and **1_*n*_** is the column-vector of ones. Using the top *K* eigenvalues and corresponding eigenvectors *U*^(*K*)^, *D*^(*K*)^, *V*^(*K*)^, from the SVD, it is known that
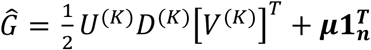 minimizes
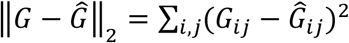 among all possible rank *K* matrices^11^, making it a good proxy for the ISAF matrix.

For a new individual s with genetic ancestry represented as *x_s_* ∈ ℝ^*k*^ in the PC (eigenvector) space of the reference panel, the ISAF for *i*-th variant can be modelled as
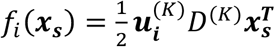 + *μ_i_*, where
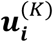 is *i*-th row of *U*^(*K*)^ and *μ_i_* is the *i*-th element of *μ*. To avoid boundary condition, we constrain
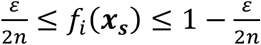 for a fixed *ε* (we used *ε* = 0.5 in our experiments). Then the overall likelihood of an individual’s genetic ancestry *x* is

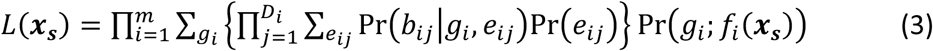

where *g_i_* represents the unobserved genotype of the sequenced sample at variant *i*. The maximum-likelihood genetic ancestry coordinates can be estimated as
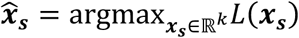 using the Nelder-Mead^12^ algorithm, starting with PC coordinates of a randomly selected individual from the reference panel. In our experiments, we always obtained consistent estimates of
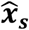 regardless of start values.

### Joint estimation of genetic ancestry and DNA contamination

Because our goal is to obtain unbiased estimates of the DNA contamination rate *α* agonistic to genetic ancestry, we propose to jointly estimate *α* and ancestry by combining the models described in the previous sections. Let ***x*_1_, *x*_2_** ∈ *R^K^* be the genetic ancestries of the intended and contaminating samples. Then the likelihood under the combined model is

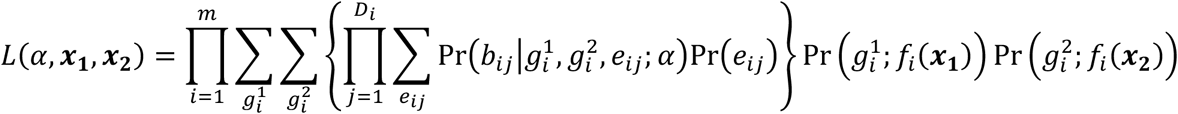

When the contamination rate *α*≈0, the parameters corresponding to ***x*_2_** do not contribute (much) to the likelihood and the estimates of ***x*_2_** may be unstable. To address this problem, we initially assume that the intended and contaminating samples are from the same population ***x*_1_** = ***x*_2_** (‘equal-ancestry’ model) and then repeat the analysis allowing for ***x*_1_** ≠ ***x*_2_** (‘unequal-ancestry’ model). The dimension of parameter space for the unequal-ancestry model is 2*k* + 1. We choose final parameter estimates between the two models based on Akaike Information Criterion (AIC)^13^.

### Evaluation on *in*-*silico* contaminated data based on 1000 Genomes project samples

We constructed *in*-*silico* contaminated DNA sequence reads using aligned low-coverage whole genome sequence reads from the 1000 Genomes phase 3 project^4^. We filtered out unmapped and mark-duplicated reads and then randomly sampled aligned sequence reads proportional to the intended contamination rates *α* ∈ {0.01,0.02,0.05,0.1,0.2}. To match the mixing proportion of sequence reads originated from intended and contaminating to be (1 – *α*): *α*, each read was sampled with probability
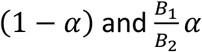 from each sample, where *B*_1_ and *B*_2_ are number of aligned bases from unique reads from intended and contaminating samples. We selected four populations, CHS (Han Chinese South), GBR (British in England and Scotland), MXL (Mexican Ancestry from Los Angeles USA), YRI (Yoruba in Ibadan, Nigeria), and arbitrarily selected 10 pairs of individuals with similar sequencing depths within the same population and across populations. To estimate genetic ancestry and/or contamination rate for these *in*-*silico* contaminated sequence reads, we used a reference panel of 938 HGDP^3^ individuals across 1,000, 10,000 and 100,000 randomly chosen SNPs (pooled MAF > 0.5%), avoiding variants masked by the 1000 Genomes Project^4^ (See Web Resource). When we compared estimated genetic ancestry with *LASER*, we used the same set of selected SNPs and sequence reads as input. For *TRACE*, we used genotypes from the phase 3 release (for 1000 Genomes) or an interim callset from the *GotCloud* software tool^14^ (for InPSYght, see next section for details) on the same SNP set.

### Experiment with real sequence data from the InPSYght study

Next, we applied our method to 500 deeply sequenced (mean depth 32x) genomes from the first two batches of the InPSYght study. For each sample, we evaluated the results from the six models: (1) the original *verifyBamID* using pooled allele frequencies; the original *verifyBamID* using (2) African, (3) East Asian, and (4) European allele frequencies; (5) the new *verifyBamID2* under the equal-ancestry model; and (6) *verifyBamID2* under the unequal-ancestry model. To calculate pooled, population-specific, and individual-specific allele frequencies, we used the 1000 Genomes phase 3 reference panel (n=2,504), randomly selecting 100,000 SNPs among the sites also polymorphic in Illumina Human Omni 2.5 array, with the same filtering criteria (MAF > 5% and 1000 Genomes mask) as above.

## Results

We assessed our new methods in the following steps. First, in the absence of contamination, we demonstrate that our estimation of genetic ancestry provides comparably accurate estimates of genetic ancestry as other state-of-art methods. Second, in the presence of contamination, we demonstrate that joint estimation of genetic ancestry and contamination substantially improves the estimation accuracy of both parameters. Third, using *in*-*silico* contaminated samples, we demonstrate that our methods robustly provide more accurate estimates than previous methods across various combinations of genetic ancestries and contamination rates. Fourth, from the analysis of deeply sequenced genomes in the InPSYght study, we demonstrate that our new methods deliver more accurate contamination estimates than the previous methods.

### New model-based methods accurately estimate genetic ancestry

In the absence of contamination, widely used methods such as *LASER* and *TRACE* are known to estimate genetic ancestry accurately. Because we propose using a new model-based approach to estimate the genetic ancestry (jointly with contamination rates), we first compared the accuracy of our new method, in the absence of contamination, with *LASER* and *TRACE.* We randomly chose 500 ethnically diverse samples from the 1000 Genomes Project low-coverage (4X) genomes, and 500 African American samples from the deeply sequenced (32x) genomes from the InPSYght project. We estimated their genetic ancestries using 100,000 SNPs from the HGDP reference panel (see Methods for details) and compared their genetic ancestry estimates obtained by LASER (using the same sequence data), and TRACE (using the hard-call genotypes). As illustrated in Figure 1A, 1C, 1E, the estimated PC coordinates of the 1000 Genomes individuals are located close to their corresponding HGDP populations across all three methods. Compared to *TRACE* and *LASER*, we observed that the estimated genetic coordinates from *verifyBamID2* were the closest to the centroid of corresponding HGDP population (Table 2) in 4 of the 5 populations (all except TSI). These results suggest that our method provides estimates at least as precise compared to those for other state-of-the-art methods.

**Figure 1.**
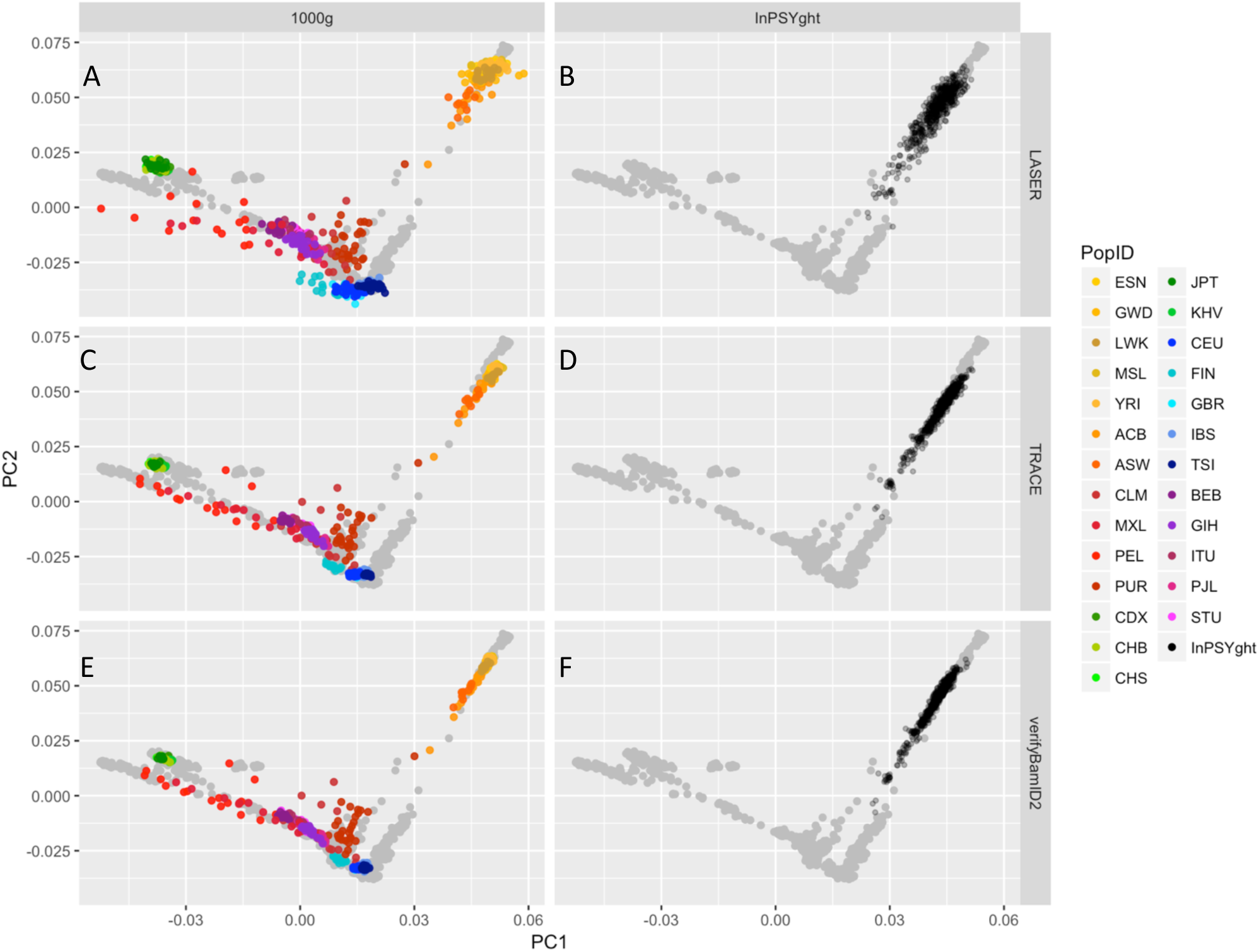
Evaluation of estimated genetic ancestry coordinates, in the absence of contamination, between *TRACE*, *LASER*, and *verifyBamID2* on samples from the 1000 Genomes low coverage genome (n=500, diverse ancestry) sequence data (A,C,E) and from the InPSYght deep genome (n=500, African Americans) sequence data (B,D,F). Panels A and B show results from *TRACE*, C and D from *LASER*, and E and F from *verifyBamID2* (assuming no contamination). Each point represents a sample and each color represents a population ancestry with the exception that grey point represents PCA coordinates of reference (HGDP) samples.

**Table 2.**
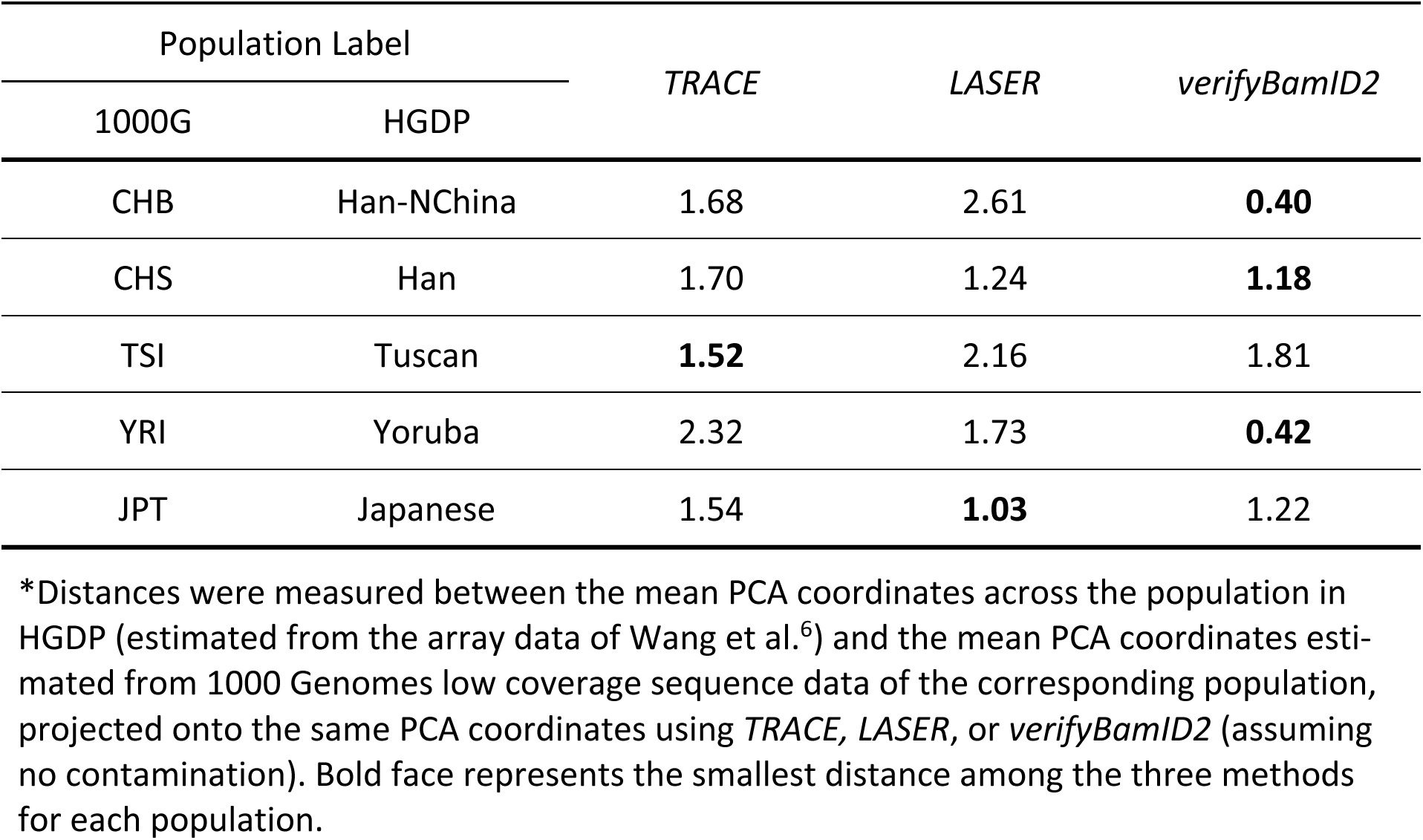
Distance between estimated PCA coordinates of HGDP and 1000G populations^∗^

| Population Label |  | *TRACE* | *LASER* | *verifyBamID2* |
| --- | --- | --- | --- | --- |
| 1000G | HGDP |  |  |  |
| CHB | Han-NChina | 1.68 | 2.61 | **0.40** |
| CHS | Han | 1.70 | 1.24 | **1.18** |
| TSI | Tuscan | **1.52** | 2.16 | 1.81 |
| YRI | Yoruba | 2.32 | 1.73 | **0.42** |
| JPT | Japanese | 1.54 | **1.03** | 1.22 |

\*Distances were measured between the mean PCA coordinates across the population in HGDP (estimated from the array data of Wang et al.<sup>6</sup>) and the mean PCA coordinates estimated from 1000 Genomes low coverage sequence data of the corresponding population, projected onto the same PCA coordinates using *TRACE*, *LASER*, or *verifyBamID2* (assuming no contamination). Bold face represents the smallest distance among the three methods for each population.

**Table 3.** Average contamination estimates for 5% contaminated samples (size n=10).

| Sample Population |  | Original Model (Fixed Allele Frequencies) |  |  |  | Equal-Ancestry Model | Unequal-Ancestry Model |
| --- | --- | --- | --- | --- | --- | --- | --- |
| Intended | Contaminating | European | East Asian | African | Pooled |  |  |
| GBR | GBR | **4.73%** | 3.19% | 2.67% | 3.76% | 4.63% | 4.63% |
| CHS | CHS | 1.90% | 4.73% | 1.25% | 2.38% | 4.73% | **4.76%** |
| YRI | YRI | 1.78% | 1.58% | **4.44%** | 2.45% | 4.40% | 4.40% |
| CHS | YRI | 3.33% | 6.91% | 2.27% | 4.10% | 6.71% | **4.81%** |
| YRI | CHS | 2.79% | 2.55% | 6.29% | 3.76% | 5.99% | **4.67%** |
| GBR | YRI | 6.13% | 4.16% | 3.60% | 5.04% | 5.90% | **4.83%** |
| YRI | GBR | 2.81% | 2.57% | 6.38% | 3.80% | 6.01% | **4.63%** |
| CHS | GBR | 2.87% | 6.33% | 1.98% | 3.55% | 6.13% | **4.83%** |
| GBR | CHS | 5.32% | 3.78% | 3.05% | 4.32% | **5.16%** | 4.67% |

Average contamination estimates of *in-silico* contaminated samples when the true contamination rate is 5%. Each mixing configuration (e.g. GBR+CHS) contains 10 samples that are constructed with 95% reads coming from the intended sample and 5% reads from the contaminating sample. The estimated contamination rates are obtained using the original version *verifyBamID* by specifying prior allele frequencies as European, East Asian, African, and Pooled, respectively. Bold represents the closest estimate to the true value of 5%.

### Genetic ancestry estimates may be confounded by DNA contamination

Next, we constructed *in*-*silico* contaminated sequenced data from the 1000 Genomes Project and estimated contamination parameters and genetic ancestries jointly. We observed that when sequences are contaminated between different continental populations, the genetic ancestry estimates in PC coordinates drift towards the contaminating population when contamination is ignored (Figure 2A) or when assuming that intended and contaminating samples originated from the same population (Figure 2B). As the contamination rate increases, drift increases.

**Figure 2.**
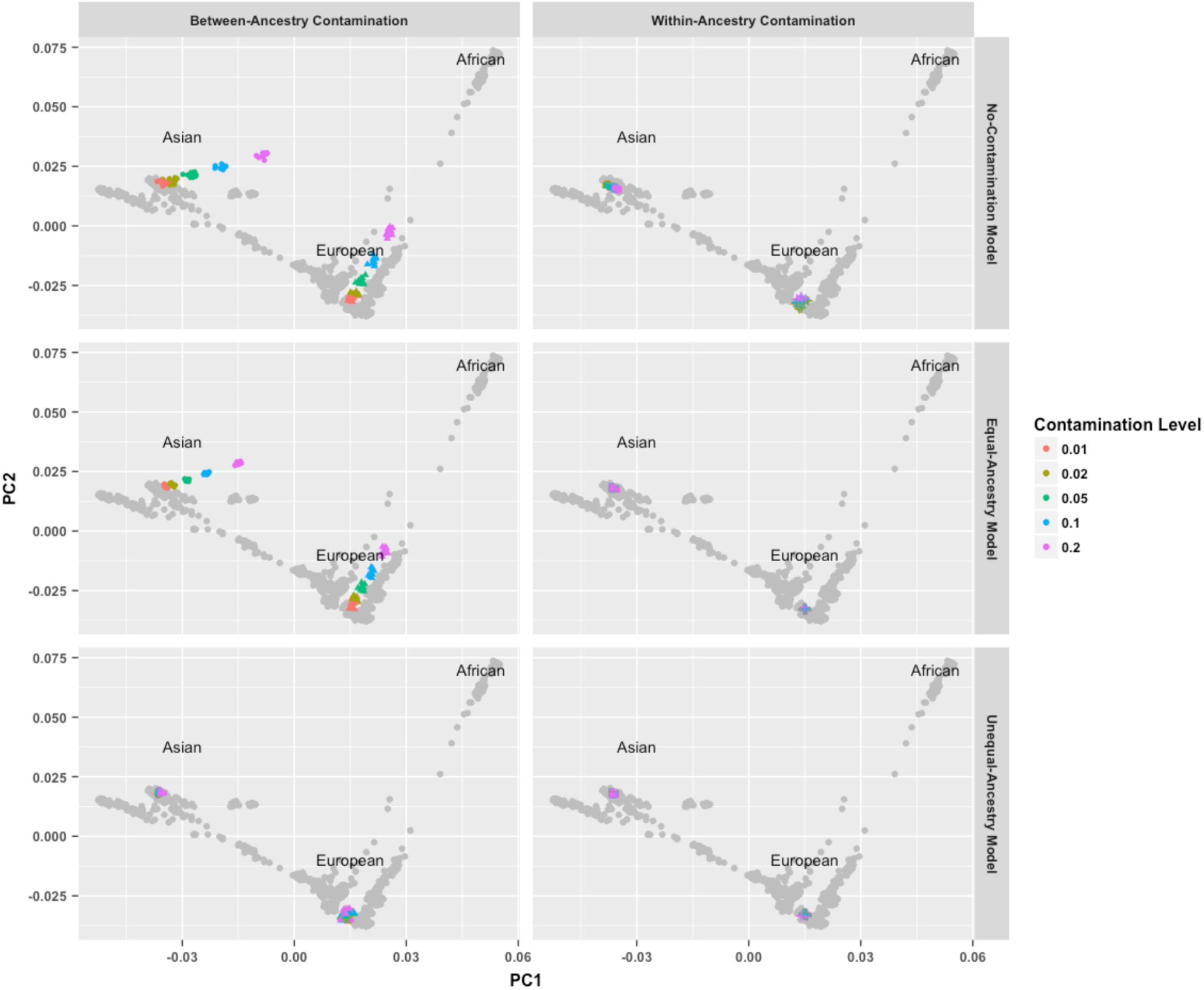
Impact of DNA sample contamination on the estimation of genetic ancestry. Each point represents a sample. Each grey point represents reference (HGDP) sample and its PCA coordinates, similar to Figure 1. Each colored point represents *in*-*silico* contaminated samples across various contamination rates and populations. In panels A, C, and E, European (GBR) and East Asian (CHS) samples are contaminated with African (YRI) samples at different contamination rates (i.e. between-ancestry contamination). In panel B, D, and F, European (GBR) and East Asian (CHS) samples are contamination with another sample in the same population (i.e. within-ancestry contamination). Different colors represent different contamination rates ranging from 1% to 20%. Upper panels (A, B) show *verifyBamID2* estimates without modelling contamination, middle panels (C, D) *verifyBamID2* estimates under the assumption that intended and contaminating populations are identical (i.e. equal-ancestry model), lower panels (E, F) *verifyBamID2* estimates under the assumption that intended and contaminating populations can be different (i.e. unequal-ancestry model).

However, when we accounted for possible differences in genetic ancestries between the two intended and contaminating samples using our new methods, PC coordinates remained similar to those for uncontaminated samples (Figure 2E), and contaminated samples constructed from individuals that belong to the same population (Figure 2B, 2D, 2F).

### Robust, accurate, ancestry-agnostic estimation of DNA contamination

Next, we evaluated the effect of genetic ancestry misspecification in estimating DNA contamination rates. We constructed contaminated samples between various combinations of populations, and compared the accuracy of estimated contamination rates using both the original methods which assume known allele frequencies and the new methods which estimate contamination rate and genetic ancestry jointly.

When contamination happens within the same population, running original methods with correct continental population allele frequencies specified provided accurate contamination estimates (Figure 3A, 3E, 3I). However, using pooled allele frequencies, which would be a default option when it is infeasible to specify population information *a priori* before sequencing, consistently underestimated contamination rates. Bias was particularly large when intended individuals were of African ancestry.

**Figure 3.**
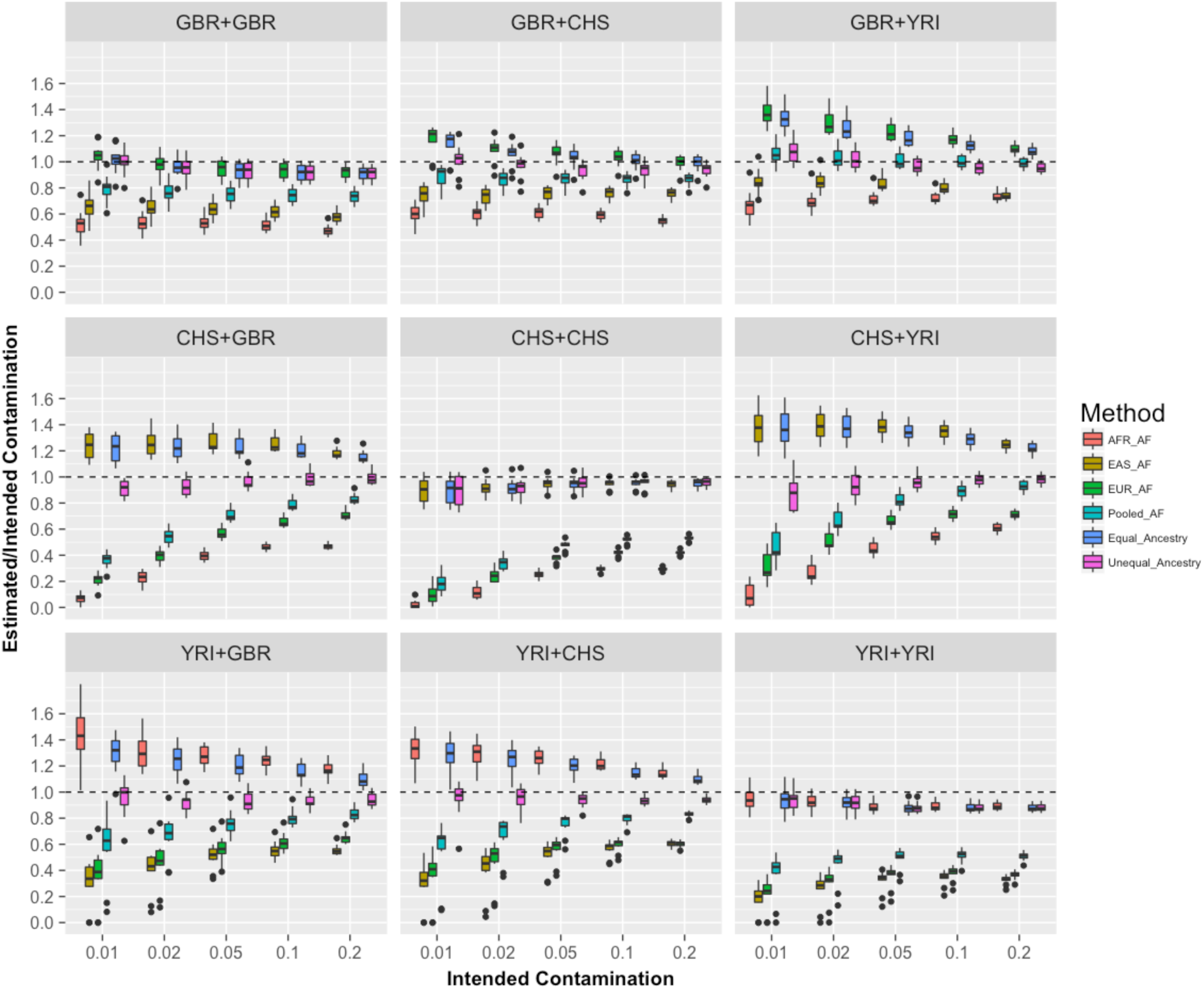
Comparison of different models to estimate contamination rates. Horizontal (x) axis shows intended contamination rate, vertical (y) axis shows the ratio of estimated to intended contamination rates. Each color represents different models to estimate contamination rates. EUR_AF, EAS_AF, and AFR_AF represent original *verifyBamID* using European, East Asian, and African allele frequencies across the continental population using the 1000 Genomes data. Pooled_AF represents the original *verifyBamID* using aggregated allele frequencies across all 2,504 individuals in the 1000 Genomes Project. Equal_Ancestry represents the *verifyBamID2* assuming that intended and contaminating samples belong to the same population. Unequal_Ancestry represents *verifyBamID2* allowing different genetic ancestry between intended and contaminating sample (recommended setting). Each panel represents different combinations of intended (row) and contaminating (column) populations, in the order of GBR, CHS, and YRI.

Specifying incorrect population allele frequencies results in even larger contamination estimation bias. For example, using African allele frequencies on East Asian samples resulted in an average estimate of 2.9% contamination for samples with contamination 10% (Table S1), implying that a large fraction of 10% contaminated samples within East Asian ancestry would not have been flagged for contamination-based exclusion at the contamination-exclusion threshold of 1-3% used by many studies e.g. the Trans-Omics Precision Medicine (TOPMed) study^15^.

Our results consistently demonstrated that the ancestry-agnostic method provides as accurate estimates as the original methods specified with correct population labels (Figure 3A, 3E, 3I, Table S1), and the estimates are substantially better than those from pooled allele frequencies or incorrectly specified allele frequencies.

When the intended and contaminating populations are different, we observed that contamination is sometimes overestimated due to increased fraction of heterozygous genotypes than expected by a given contamination rate under single population model. Our method based on unequal-ancestry model outperforms all the other methods in terms of overall bias and Mean Squared Error(MSE) (Figure 3, Table S4), correcting for both upward and downward biases in various ancestry combinations. For example, the relative deviation of estimated to intended contamination rate (i.e. 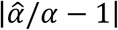) is reduced by 80% (73-88%) compared to the original *verifyBamID* with various population allele frequencies, suggesting reduced bias. MSE is also reduced by 92% (86-97%). This robustness reflects the ability to incorporate differences in population allele frequencies between intended and contaminating individuals (Figure 3B, 3C, 3D, 3F, 3G, 3H, Table S1).

We also examined the accuracy of our methods for admixed populations by performing a similar experiment using the Mexican population (MXL) and obtained consistent results (Supplementary Table S2).

### Results with deep whole genome sequence data from the InPSYght study

Next, we applied our methods to 500 African American samples from the InPSYght study (see Methods). Consistent with the results from our *in silico* contamination studies, we observed that the average contamination rate was 1.1-fold higher with newer method (0.36% for unequal-ancestry, 0.37% for equal-ancestry) compared to the original method with pooled allele frequency (0.33%) (Figure 4). The number of samples with estimated contamination rate >1% increased from 16 (original method with pooled allele frequency) to 21 (unequal-ancestry method) or 23 (unequal-ancestry method), suggesting our new method more rigorously screens for contaminated samples.

**Figure 4.**
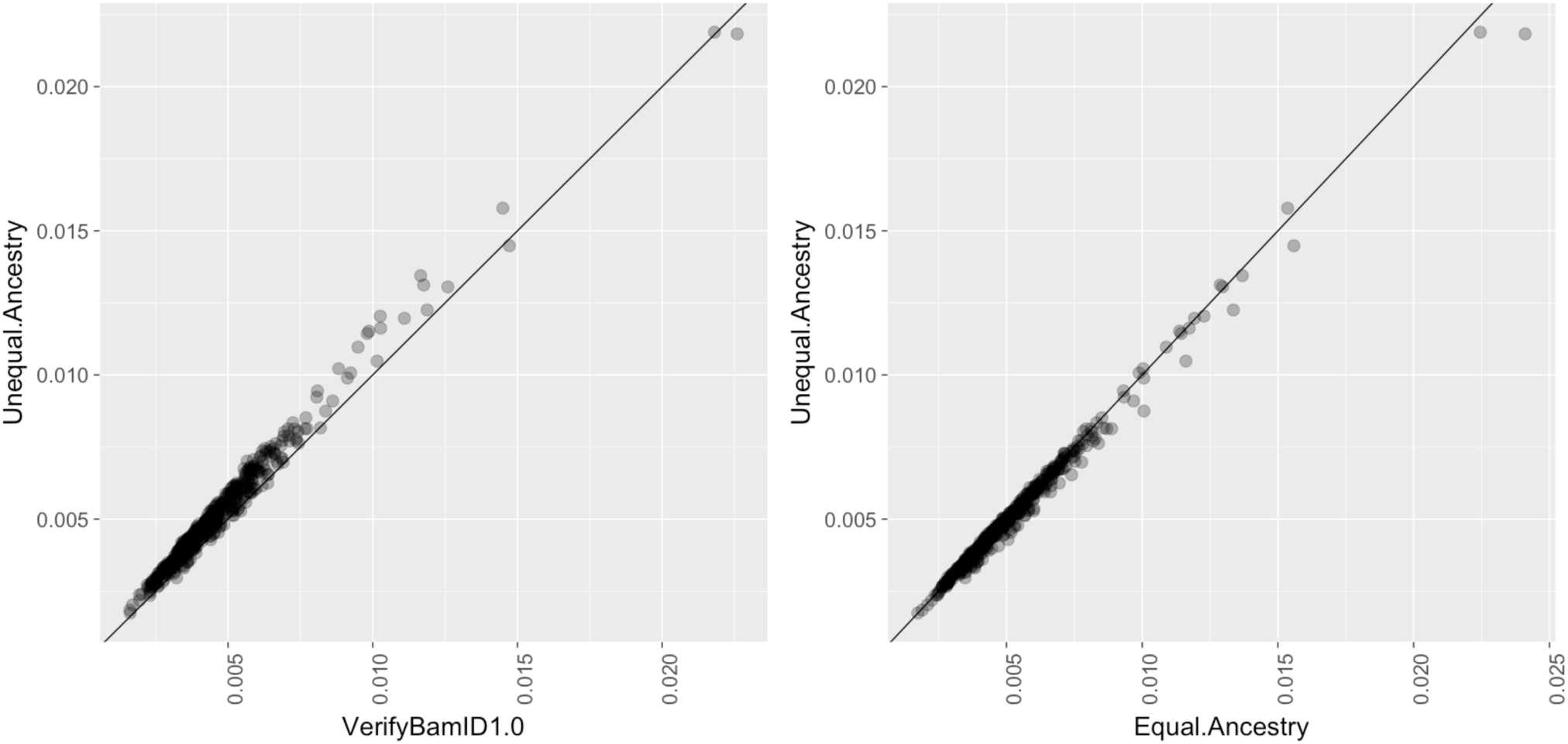
Comparison of contamination estimation between using *verifyBamID* and *verifyBamID2* on 500 InPSYght samples. All subjects are African Americans. Each dot represents the pair of contamination rate estimates using different methods. The left panel shows the estimated contamination rates of the original *verifyBamID* with pooled allele frequencies, which is the default setting of *verifyBamID* in x-axis. Y-axis shows *verifyBamID2* with unequal-ancestry model (y-axis). Each point represents a sequenced subject. The right panel compares the estimated contamination rates between two models (unequal-ancestry vs. equal-ancestry) of *verifyBamID2* on the same dataset.

All 500 deeply sequenced genomes in InPSYght study are reported to be African Americans, and indeed the estimated PC coordinates for all 500 individuals under all three methods lie between European and African samples. Compared to other methods to estimate genetic ancestry, our estimates resulted in tighter clustering along the European-African segment than *LASER*, and similarly tight clustering to *TRACE* (Figure 1B, 1D, 1F). For example, the correlation coefficient between the PC1 and PC2 coordinates were 0.927 for *LASER*, 0.981 for *TRACE*, and 0.985 for *verifyBamID2*, corroborating that *verifyBamID2* results in more precise estimate of African ancestry along the European-African segment in PC coordinates.

### Impact of number of markers on accuracy, computational cost, and memory requirements

As we have shown previously^2^, there are trade-offs between computation cost and accuracy of contamination estimates. Using as many as 100,000 variants results in accurately estimated intended contamination rate. For example, MSE of relative deviation (i.e. 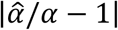) was 0.02, 0.01, 0.01 when the intended contamination was 1%, 2%, and 5%, respecitvely. When we use 10,000 variants, the MSEs modestly increased to 0.11, 0.04, and 0.01, respectively. When we use only 1,000 variants, MSEs further increased to 0.69, 0.25, 0.11, suggesting that the estimates may not be precise for low contamination rate when using only 1,000 variants. (Supplementary Table S3).

We also evaluated the computational cost and memory consumption of *verifyBamID2* on whole genome sequence data with various coverages. For the BAM files from the 1000 Genomes whole genome sequence data (4.3-5.1x coverage), the average wall-clock running time was 5.5 minutes with a single thread and peak memory consumption was 505 MB when using 10,000 markers in a server with Xeon 2.27GHz processor. When using 100,000 markers, the average wall-clock running time was 20.5 minutes with a single thread and 8.0 minutes with four threads, and peak memory consumption was 528 MB.

For deep genome data from the InPSYght study (31x coverage) stored in CRAM format, the average wall-clock time was 17.3 minutes and peak memory consumption was 514 MB when using 10,000 markers. For 100,000 markers the average wall-clock time was 155.6 minutes (single thread) or 96 minutes (four threads) and peak memory consumption was 548 MB.

## Discussion

Contamination detection is an essential step in the sequence analysis process that has important effects on following downstream analyses. Early and accurate estimation of DNA contamination can prevent wasted effort, time, and money by identifying the problems early on before too many samples are sequenced using contamination-prone protocols. Our previous method enabled such a timely contamination detection from sequence data and population allele frequencies at known variant sites, without requiring independent SNP genotype data. Our new method maintains these advantages, and in addition provide three more. First, because our joint analysis method is agnostic to genetic ancestry, it eliminates sample-to-sample variation in the parameter settings for the contamination checking procedure, simplifying the sequence analysis pipeline. Second, it provides more robust contamination estimates against potentially misspecified population allele frequency of the intended (or contaminating) samples when relying on the reported ancestry information. Third, it provides accurate estimates of genetic ancestries for both intended and contaminating samples. This enables additional sanity checking of the sequence data, such as determining whether a sequenced sample matches its expected (participant-reported) ancestry. It also facilitates incorporating ancestry information in the variant calling and downstream analysis, and allows us to track the source of contamination more precisely when contamination occurs.

Our method can be used not only to detect and estimate contamination, but also to estimate genetic ancestry from sequence data. Relatively few methods, such as *LASER*^5,6^ and *bammds*^16^, exist for estimating genetic ancestry from sequence data while several methods have been developed for array-based genotypes, such as *EIGENSOFT*^17^, *FRAPPE*^18^, *ADMIXTURE*^19^, and *TRACE*^6^. We have demonstrated that our method provides ancestry estimates as or more accurate than *LASER*, particularly when the sequenced samples are contaminated between different ancestries.

By jointly estimating genetic ancestry and contamination, we are able to accurately estimate contamination without requiring ancestry information *a priori.* Since obtaining population allele frequency information may be infeasible or even impossible at the time of sequencing, it is important to highlight that our ancestry-agnostic approach provides more timely and accurate feedback to the sequencing facilities. Our ancestry-agnostic approach also simplifies the sequence analysis pipeline, because the same input arguments can be applied across all samples regardless of their genetic ancestry

The key idea of using individual-specific allele frequencies (ISAF) to account for population structure in genetic analysis has been suggested previously in the context of characterizing population structure or identifying highly differentiated variants across populations^8,9^. To the best our knowledge, our method describes the first likelihood-based model utilizing ISAF to represent high throughput sequence reads under population structure and/or contamination. While previous studies proposed logistic models as alternative to linear model^8,9^, we used linear models (bounded by minimum and maximum value) between allele frequencies and population structure represented by Singular Value Decomposition (SVD) on the genotype matrix. We made this choice because the logistic model is computationally more intensive, and the linear model is accurate for the common variants we use, as demonstrated by the previous studies^9^.

Because we use Nelder-Mead optimization for maximum likelihood estimation, it is possible that the estimates do not converge to the global maximum, especially when many principal components are used. We observed that estimating the full unequal-ancestry model parameters sometimes does fail to converge especially when there is little or no contamination, due to the limited identifiability of the genetic ancestry of contaminating samples in this situation. Starting by estimating contamination rate and shared genetic ancestry parameters using the equal-ancestry model, and using those estimates as start values for the unequal-ancestry model to allow different ancestries between the intended and contaminating samples dramatically improved convergence; in fact, the method converged to consistent estimates across multiple starting points within 1,000 iterations in all our benchmark cases, in both real and *in*-*silico* contaminated data. When the contamination rate is extremely small (e.g. <0.1%), estimation of genetic ancestry of contaminating samples can still be challenging. We allow unequal ancestries between intended and contaminating samples only when the likelihood substantially improves beyond AIC threshold between equal ancestry and unequal ancestry models. This procedure effectively removed all outlier estimates of genetic ancestries of contaminating samples in our experiments.

There are other possible useful extensions to our joint contamination and estimation method. i We are extending these methods to detect and estimate contamination for RNA-seq and other i epigenomic sequence data. The same model has potential applications in other areas, such as cancer single cell transcriptomics^20^.

We expect that our new *verifyBamID2* software will facilitate more accurate, convenient, and timely quality control of sequence genomes. Our software tool is publicly available at http://github.com/Griffan/verifyBamID. Our GitHub repository provides reference files that can be used as test input for our methods. These files contain key input files required for *verifyBamID2*, including variant loadings, supporting various genome builds (GRCh37 and GRCh38), and various numbers of variants.

## Web Resources

1000 genomes project genome mask file: (ftp://ftp.1000genomes.ebi.ac.uk/vol1/ftp/release/20130502/supporting/accessible_genome_masks/StrictMask/)

## Acknowledgements

This work was supported by NIH grants HG009976 (to M.F. and M.B.), HL137182 (to H.M.K. and F.Z), HG007022 (to G.R.A), MH105653 (to M.B, InPSYght Consortium, and H.M.K.)

---

**Supplementary Table S1:** Mean estimated contamination rates of *in*-*silico* contaminated population across different intended contamination rate, populations of intended and contaminating samples, and the estimation methods.

| Population |  | Intended<br>% Contam. | Equal-Ancestry<br>(VB2) | Unequal-Ancestry<br>(VB2) | Pooled<br>AF<br>(VB1) | EUR<br>AF<br>(VB1) | EAS<br>AF<br>(VB1) | AFR<br>AF<br>(VB1) |
| --- | --- | --- | --- | --- | --- | --- | --- | --- |
| Intended | Contam. |  |  |  |  |  |  |  |
| GBR | GBR | 1% | 1.0% | 1.0% | 0.8% | 1.0% | 0.6% | 0.5% |
|  |  | 2% | 1.9% | 1.9% | 1.5% | 2.0% | 1.3% | 1.1% |
|  |  | 5% | 4.6% | 4.6% | 3.8% | 4.7% | 3.2% | 2.7% |
|  |  | 10% | 9.2% | 9.2% | 7.4% | 9.4% | 6.2% | 5.2% |
|  |  | 20% | 18.3% | 18.3% | 14.7% | 18.5% | 11.6% | 9.5% |
| GBR | CHS | 1% | 1.1% | 1.0% | 0.9% | 1.2% | 0.7% | 0.6% |
|  |  | 2% | 2.1% | 1.9% | 1.7% | 2.2% | 1.5% | 1.2% |
|  |  | 5% | 5.2% | 4.7% | 4.3% | 5.3% | 3.8% | 3.1% |
|  |  | 10% | 10.1% | 9.4% | 8.6% | 10.4% | 7.6% | 5.9% |
|  |  | 20% | 19.8% | 18.7% | 17.3% | 19.9% | 15.1% | 10.9% |
| GBR | YRI | 1% | 1.3% | 1.1% | 1.1% | 1.4% | 0.8% | 0.7% |
|  |  | 2% | 2.5% | 2.0% | 2.1% | 2.6% | 1.7% | 1.4% |
|  |  | 5% | 5.9% | 4.8% | 5.0% | 6.1% | 4.2% | 3.6% |
|  |  | 10% | 11.3% | 9.5% | 10.0% | 11.7% | 8.0% | 7.3% |
|  |  | 20% | 21.6% | 19.1% | 19.7% | 22.0% | 14.8% | 14.6% |
| CHS | GBR | 1% | 1.2% | 0.9% | 0.4% | 0.2% | 1.2% | 0.1% |
|  |  | 2% | 2.5% | 1.8% | 1.1% | 0.8% | 2.5% | 0.5% |
|  |  | 5% | 6.1% | 4.8% | 3.6% | 2.9% | 6.3% | 2.0% |
|  |  | 10% | 12.0% | 9.9% | 7.9% | 6.6% | 12.5% | 4.6% |
|  |  | 20% | 23.0% | 19.8% | 16.6% | 14.2% | 23.6% | 9.4% |
| CHS | CHS | 1% | 0.9% | 0.9% | 0.2% | 0.1% | 0.9% | 0.0% |
|  |  | 2% | 1.8% | 1.8% | 0.7% | 0.5% | 1.8% | 0.2% |
|  |  | 5% | 4.7% | 4.8% | 2.4% | 1.9% | 4.7% | 1.3% |
|  |  | 10% | 9.5% | 9.5% | 5.2% | 4.2% | 9.5% | 2.9% |
|  |  | 20% | 19.1% | 19.1% | 10.6% | 8.4% | 18.9% | 5.9% |
| CHS | YRI | 1% | 1.4% | 0.9% | 0.5% | 0.3% | 1.4% | 0.1% |
|  |  | 2% | 2.8% | 1.9% | 1.3% | 1.0% | 2.8% | 0.5% |
|  |  | 5% | 6.7% | 4.8% | 4.1% | 3.3% | 6.9% | 2.3% |
|  |  | 10% | 12.9% | 9.8% | 8.9% | 7.2% | 13.5% | 5.4% |
|  |  | 20% | 24.3% | 19.6% | 18.6% | 14.2% | 24.9% | 12.2% |
| YRI | GBR | 1% | 1.3% | 1.0% | 0.6% | 0.4% | 0.4% | 1.4% |
|  |  | 2% | 2.5% | 1.9% | 1.3% | 0.9% | 0.8% | 2.6% |
|  |  | 5% | 6.0% | 4.6% | 3.8% | 2.8% | 2.6% | 6.4% |
|  |  | 10% | 11.5% | 9.3% | 8.1% | 6.2% | 5.6% | 12.5% |
|  |  | 20% | 21.9% | 18.8% | 16.7% | 13.0% | 11.1% | 23.5% |
| YRI | CHS | 1% | 1.3% | 0.9% | 0.5% | 0.4% | 0.3% | 1.3% |
|  |  | 2% | 2.5% | 1.9% | 1.3% | 0.9% | 0.8% | 2.6% |
|  |  | 5% | 6.0% | 4.7% | 3.8% | 2.8% | 2.5% | 6.3% |
|  |  | 10% | 11.5% | 9.3% | 7.9% | 5.9% | 5.6% | 12.2% |
|  |  | 20% | 22.0% | 18.8% | 16.6% | 12.0% | 12.1% | 22.9% |
| YRI | YRI | 1% | 0.9% | 0.9% | 0.4% | 0.2% | 0.2% | 0.9% |
|  |  | 2% | 1.8% | 1.8% | 0.9% | 0.6% | 0.5% | 1.8% |
|  |  | 5% | 4.4% | 4.4% | 2.4% | 1.8% | 1.6% | 4.4% |
|  |  | 10% | 8.8% | 8.8% | 5.1% | 3.8% | 3.4% | 8.9% |
|  |  | 20% | 17.6% | 17.6% | 10.1% | 7.3% | 6.6% | 17.9% |

Equal-Ancestry Model: Estimate from *verifyBamID2* assuming intended and contaminating samples have the same genetic ancestry (in PC coordinates)
Unequal-Ancestry Model: Estimate from *verifyBamID2* allowing intended and contaminating samples to have different genetic ancestry
Pooled AF: Estimate from original *verifyBamID* using allele frequency across all 1000 Genomes phase 3 samples
EUR AF: Estimate from original *verifyBamID* using allele frequency across European subset of 1000 Genomes phase 3 samples
EAS AF: Estimate from original *verifyBamID* using allele frequency across East Asian subset of 1000 Genomes phase 3 samples
AFR AF: Estimate from original *verifyBamID* using allele frequency across African subset of 1000 Genomes phase 3 samples

**Supplementary Table S2:** Average of estimated contamination rates across 10 *in*-*silico* contaminated samples from Mexican population under different models. Results are similar as Europeans, except that unequal-ancestry model slightly reduces estimated contamination rate from equal-ancestry model, unlike GBR.

| Intended %<br>Contamination | Equal-Ancestry<br>(VB2) | Unequal-Ancestry<br>(VB2) | Pooled<br>AF<br>(VB1) | EUR<br>AF<br>(VB1) | EAS<br>AF<br>(VB1) | AFR<br>AF<br>(VB1) |
| --- | --- | --- | --- | --- | --- | --- |
| 1% | 1.1% | 1.0% | 0.8% | 1.0% | 0.6% | 0.3% |
| 2% | 2.1% | 2.1% | 1.6% | 2.0% | 1.4% | 0.9% |
| 5% | 4.8% | 4.8% | 3.9% | 4.6% | 3.5% | 2.5% |
| 10% | 9.3% | 9.2% | 7.8% | 8.8% | 6.8% | 4.9% |
| 20% | 18.5% | 18.3% | 15.4% | 17.0% | 13.0% | 9.4% |

**Supplementary Table S3:**
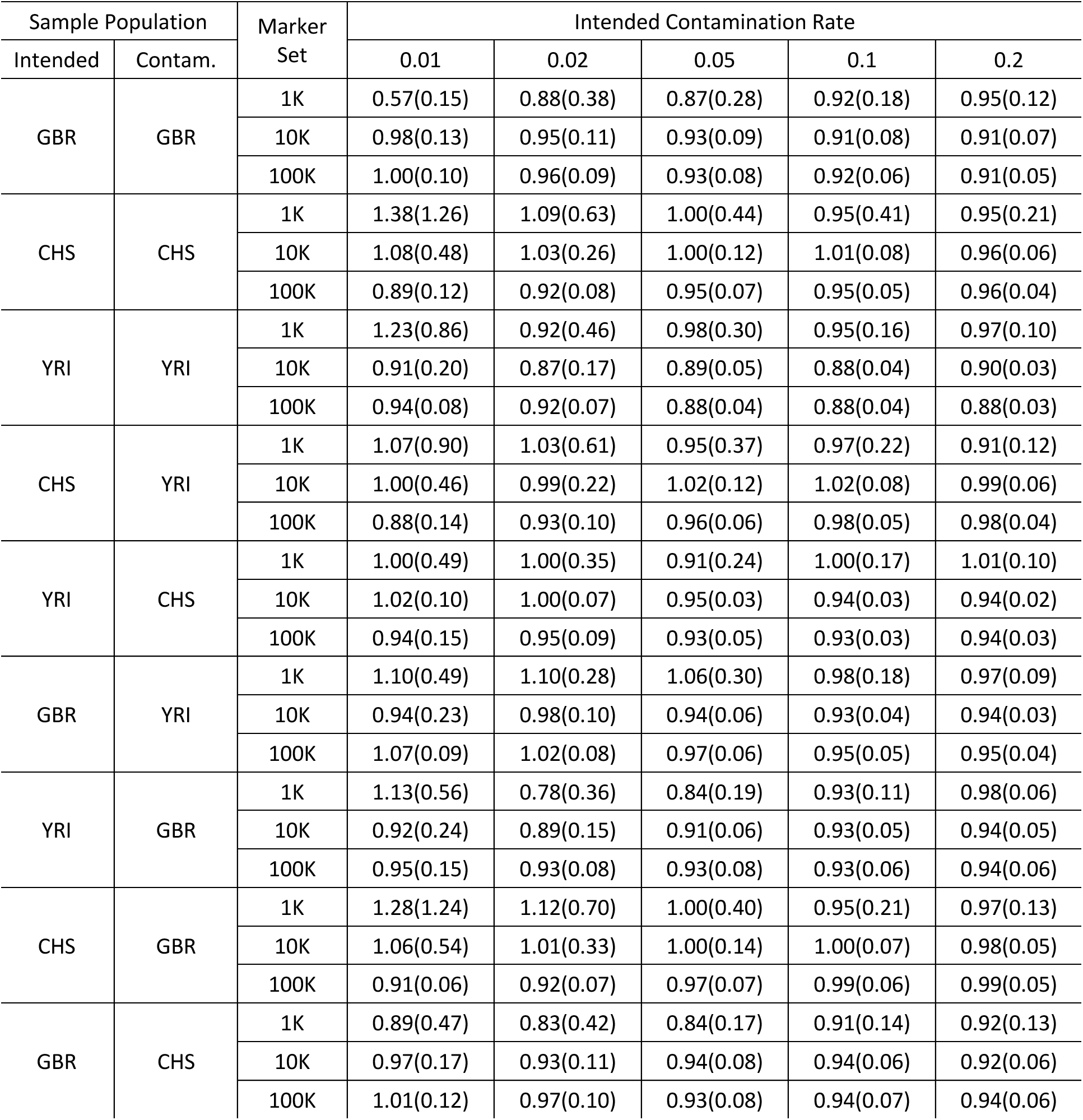
Comparison of mean contamination rate ratio (Estimated/Intended) using different size of marker set (under Unequal-Ancestry Model). The Numbers in parenthesis represent standard deviation.

| Sample Population |  | Marker Set | Intended Contamination Rate |  |  |  |  |
| --- | --- | --- | --- | --- | --- | --- | --- |
| Intended | Contam. |  | 0.01 | 0.02 | 0.05 | 0.1 | 0.2 |
| GBR | GBR | 1K | 0.57(0.15) | 0.88(0.38) | 0.87(0.28) | 0.92(0.18) | 0.95(0.12) |
|  |  | 10K | 0.98(0.13) | 0.95(0.11) | 0.93(0.09) | 0.91(0.08) | 0.91(0.07) |
|  |  | 100K | 1.00(0.10) | 0.96(0.09) | 0.93(0.08) | 0.92(0.06) | 0.91(0.05) |
| CHS | CHS | 1K | 1.38(1.26) | 1.09(0.63) | 1.00(0.44) | 0.95(0.41) | 0.95(0.21) |
|  |  | 10K | 1.08(0.48) | 1.03(0.26) | 1.00(0.12) | 1.01(0.08) | 0.96(0.06) |
|  |  | 100K | 0.89(0.12) | 0.92(0.08) | 0.95(0.07) | 0.95(0.05) | 0.96(0.04) |
| YRI | YRI | 1K | 1.23(0.86) | 0.92(0.46) | 0.98(0.30) | 0.95(0.16) | 0.97(0.10) |
|  |  | 10K | 0.91(0.20) | 0.87(0.17) | 0.89(0.05) | 0.88(0.04) | 0.90(0.03) |
|  |  | 100K | 0.94(0.08) | 0.92(0.07) | 0.88(0.04) | 0.88(0.04) | 0.88(0.03) |
| CHS | YRI | 1K | 1.07(0.90) | 1.03(0.61) | 0.95(0.37) | 0.97(0.22) | 0.91(0.12) |
|  |  | 10K | 1.00(0.46) | 0.99(0.22) | 1.02(0.12) | 1.02(0.08) | 0.99(0.06) |
|  |  | 100K | 0.88(0.14) | 0.93(0.10) | 0.96(0.06) | 0.98(0.05) | 0.98(0.04) |
| YRI | CHS | 1K | 1.00(0.49) | 1.00(0.35) | 0.91(0.24) | 1.00(0.17) | 1.01(0.10) |
|  |  | 10K | 1.02(0.10) | 1.00(0.07) | 0.95(0.03) | 0.94(0.03) | 0.94(0.02) |
|  |  | 100K | 0.94(0.15) | 0.95(0.09) | 0.93(0.05) | 0.93(0.03) | 0.94(0.03) |
| GBR | YRI | 1K | 1.10(0.49) | 1.10(0.28) | 1.06(0.30) | 0.98(0.18) | 0.97(0.09) |
|  |  | 10K | 0.94(0.23) | 0.98(0.10) | 0.94(0.06) | 0.93(0.04) | 0.94(0.03) |
|  |  | 100K | 1.07(0.09) | 1.02(0.08) | 0.97(0.06) | 0.95(0.05) | 0.95(0.04) |
| YRI | GBR | 1K | 1.13(0.56) | 0.78(0.36) | 0.84(0.19) | 0.93(0.11) | 0.98(0.06) |
|  |  | 10K | 0.92(0.24) | 0.89(0.15) | 0.91(0.06) | 0.93(0.05) | 0.94(0.05) |
|  |  | 100K | 0.95(0.15) | 0.93(0.08) | 0.93(0.08) | 0.93(0.06) | 0.94(0.06) |
| CHS | GBR | 1K | 1.28(1.24) | 1.12(0.70) | 1.00(0.40) | 0.95(0.21) | 0.97(0.13) |
|  |  | 10K | 1.06(0.54) | 1.01(0.33) | 1.00(0.14) | 1.00(0.07) | 0.98(0.05) |
|  |  | 100K | 0.91(0.06) | 0.92(0.07) | 0.97(0.07) | 0.99(0.06) | 0.99(0.05) |
| GBR | CHS | 1K | 0.89(0.47) | 0.83(0.42) | 0.84(0.17) | 0.91(0.14) | 0.92(0.13) |
|  |  | 10K | 0.97(0.17) | 0.93(0.11) | 0.94(0.08) | 0.94(0.06) | 0.92(0.06) |
|  |  | 100K | 1.01(0.12) | 0.97(0.10) | 0.93(0.08) | 0.94(0.07) | 0.94(0.06) |

**Supplementary Table S4:** A full table summarizing the contamination rate ratio (Estimated/Intended) across various simulation parameters, populations, and estimation methods shown in Figure 3. 100K marker sets were used.

| Sample Population |  | Method | Allele Frequencies | Intended<br>% Contam | Mean | SD | MSE |
| --- | --- | --- | --- | --- | --- | --- | --- |
| In-<br>tended | Contam. |  |  |  |  |  |  |
| GBR | GBR | VB1 | AFR | 1% | 0.52 | 0.11 | 0.242 |
| GBR | GBR | VB1 | AFR | 2% | 0.53 | 0.09 | 0.223 |
| GBR | GBR | VB1 | AFR | 5% | 0.53 | 0.06 | 0.221 |
| GBR | GBR | VB1 | AFR | 10% | 0.52 | 0.05 | 0.237 |
| GBR | GBR | VB1 | AFR | 20% | 0.48 | 0.04 | 0.276 |
| GBR | GBR | VB1 | EUR | 1% | 1.04 | 0.10 | 0.012 |
| GBR | GBR | VB1 | EUR | 2% | 0.98 | 0.09 | 0.007 |
| GBR | GBR | VB1 | EUR | 5% | 0.95 | 0.07 | 0.008 |
| GBR | GBR | VB1 | EUR | 10% | 0.94 | 0.06 | 0.008 |
| GBR | GBR | VB1 | EUR | 20% | 0.92 | 0.05 | 0.008 |
| GBR | GBR | VB1 | EAS | 1% | 0.65 | 0.11 | 0.136 |
| GBR | GBR | VB1 | EAS | 2% | 0.65 | 0.09 | 0.132 |
| GBR | GBR | VB1 | EAS | 5% | 0.64 | 0.06 | 0.135 |
| GBR | GBR | VB1 | EAS | 10% | 0.62 | 0.05 | 0.148 |
| GBR | GBR | VB1 | EAS | 20% | 0.58 | 0.05 | 0.179 |
| GBR | GBR | VB1 | Pooled | 1% | 0.79 | 0.11 | 0.055 |
| GBR | GBR | VB1 | Pooled | 2% | 0.77 | 0.08 | 0.060 |
| GBR | GBR | VB1 | Pooled | 5% | 0.75 | 0.07 | 0.066 |
| GBR | GBR | VB1 | Pooled | 10% | 0.74 | 0.06 | 0.069 |
| GBR | GBR | VB1 | Pooled | 20% | 0.73 | 0.05 | 0.073 |
| GBR | GBR | VB2 | ISAF (Equal-Ancestry) | 1% | 1.02 | 0.11 | 0.010 |
| GBR | GBR | VB2 | ISAF (Equal-Ancestry) | 2% | 0.96 | 0.09 | 0.009 |
| GBR | GBR | VB2 | ISAF (Equal -Ancestry) | 5% | 0.93 | 0.07 | 0.010 |
| GBR | GBR | VB2 | ISAF (Equal -Ancestry) | 10% | 0.92 | 0.06 | 0.010 |
| GBR | GBR | VB2 | ISAF (Equal -Ancestry) | 20% | 0.91 | 0.05 | 0.010 |
| GBR | GBR | VB2 | ISAF (Unequal-Ancestry) | 1% | 1.00 | 0.10 | 0.009 |
| GBR | GBR | VB2 | ISAF (Unequal-Ancestry) | 2% | 0.96 | 0.09 | 0.009 |
| GBR | GBR | VB2 | ISAF (Unequal-Ancestry) | 5% | 0.93 | 0.08 | 0.011 |
| GBR | GBR | VB2 | ISAF (Unequal-Ancestry) | 10% | 0.92 | 0.06 | 0.010 |
| GBR | GBR | VB2 | ISAF (Unequal-Ancestry) | 20% | 0.91 | 0.05 | 0.010 |
| GBR | CHS | VB1 | AFR | 1% | 0.59 | 0.08 | 0.172 |
| GBR | CHS | VB1 | AFR | 2% | 0.60 | 0.06 | 0.162 |
| GBR | CHS | VB1 | AFR | 5% | 0.61 | 0.05 | 0.154 |
| GBR | CHS | VB1 | AFR | 10% | 0.59 | 0.04 | 0.169 |
| GBR | CHS | VB1 | AFR | 20% | 0.55 | 0.03 | 0.206 |
| GBR | CHS | VB1 | EUR | 1% | 1.17 | 0.11 | 0.039 |
| GBR | CHS | VB1 | EUR | 2% | 1.09 | 0.10 | 0.016 |
| GBR | CHS | VB1 | EUR | 5% | 1.06 | 0.08 | 0.010 |
| GBR | CHS | VB1 | EUR | 10% | 1.04 | 0.07 | 0.006 |
| GBR | CHS | VB1 | EUR | 20% | 0.99 | 0.06 | 0.003 |
| GBR | CHS | VB1 | EAS | 1% | 0.74 | 0.09 | 0.074 |
| GBR | CHS | VB1 | EAS | 2% | 0.74 | 0.07 | 0.072 |
| GBR | CHS | VB1 | EAS | 5% | 0.76 | 0.06 | 0.063 |
| GBR | CHS | VB1 | EAS | 10% | 0.76 | 0.05 | 0.061 |
| GBR | CHS | VB1 | EAS | 20% | 0.75 | 0.04 | 0.062 |
| GBR | CHS | VB1 | Pooled | 1% | 0.89 | 0.09 | 0.019 |
| GBR | CHS | VB1 | Pooled | 2% | 0.86 | 0.07 | 0.025 |
| GBR | CHS | VB1 | Pooled | 5% | 0.86 | 0.06 | 0.022 |
| GBR | CHS | VB1 | Pooled | 10% | 0.86 | 0.06 | 0.022 |
| GBR | CHS | VB1 | Pooled | 20% | 0.86 | 0.05 | 0.021 |
| GBR | CHS | VB2 | ISAF (Equal-Ancestry) | 1% | 1.13 | 0.11 | 0.028 |
| GBR | CHS | VB2 | ISAF (Equal-Ancestry) | 2% | 1.06 | 0.09 | 0.011 |
| GBR | CHS | VB2 | ISAF (Equal -Ancestry) | 5% | 1.03 | 0.08 | 0.007 |
| GBR | CHS | VB2 | ISAF (Equal -Ancestry) | 10% | 1.01 | 0.07 | 0.004 |
| GBR | CHS | VB2 | ISAF (Equal -Ancestry) | 20% | 0.99 | 0.06 | 0.004 |
| GBR | CHS | VB2 | ISAF (Unequal-Ancestry) | 1% | 1.01 | 0.12 | 0.012 |
| GBR | CHS | VB2 | ISAF (Unequal-Ancestry) | 2% | 0.97 | 0.10 | 0.010 |
| GBR | CHS | VB2 | ISAF (Unequal-Ancestry) | 5% | 0.93 | 0.08 | 0.010 |
| GBR | CHS | VB2 | ISAF (Unequal-Ancestry) | 10% | 0.94 | 0.07 | 0.008 |
| GBR | CHS | VB2 | ISAF (Unequal-Ancestry) | 20% | 0.94 | 0.06 | 0.007 |
| GBR | YRI | VB1 | AFR | 1% | 0.67 | 0.11 | 0.119 |
| GBR | YRI | VB1 | AFR | 2% | 0.70 | 0.09 | 0.096 |
| GBR | YRI | VB1 | AFR | 5% | 0.72 | 0.06 | 0.082 |
| GBR | YRI | VB1 | AFR | 10% | 0.73 | 0.05 | 0.077 |
| GBR | YRI | VB1 | AFR | 20% | 0.73 | 0.04 | 0.074 |
| GBR | YRI | VB1 | EUR | 1% | 1.38 | 0.10 | 0.150 |
| GBR | YRI | VB1 | EUR | 2% | 1.30 | 0.09 | 0.097 |
| GBR | YRI | VB1 | EUR | 5% | 1.23 | 0.07 | 0.055 |
| GBR | YRI | VB1 | EUR | 10% | 1.17 | 0.05 | 0.032 |
| GBR | YRI | VB1 | EUR | 20% | 1.10 | 0.04 | 0.011 |
| GBR | YRI | VB1 | EAS | 1% | 0.85 | 0.10 | 0.032 |
| GBR | YRI | VB1 | EAS | 2% | 0.85 | 0.08 | 0.028 |
| GBR | YRI | VB1 | EAS | 5% | 0.83 | 0.06 | 0.031 |
| GBR | YRI | VB1 | EAS | 10% | 0.80 | 0.04 | 0.042 |
| GBR | YRI | VB1 | EAS | 20% | 0.74 | 0.03 | 0.069 |
| GBR | YRI | VB1 | Pooled | 1% | 1.05 | 0.09 | 0.010 |
| GBR | YRI | VB1 | Pooled | 2% | 1.03 | 0.08 | 0.007 |
| GBR | YRI | VB1 | Pooled | 5% | 1.01 | 0.06 | 0.003 |
| GBR | YRI | VB1 | Pooled | 10% | 1.00 | 0.05 | 0.002 |
| GBR | YRI | VB1 | Pooled | 20% | 0.99 | 0.04 | 0.002 |
| GBR | YRI | VB2 | ISAF (Equal-Ancestry) | 1% | 1.33 | 0.09 | 0.118 |
| GBR | YRI | VB2 | ISAF (Equal-Ancestry) | 2% | 1.26 | 0.09 | 0.074 |
| GBR | YRI | VB2 | ISAF (Equal -Ancestry) | 5% | 1.18 | 0.06 | 0.036 |
| GBR | YRI | VB2 | ISAF (Equal -Ancestry) | 10% | 1.13 | 0.05 | 0.018 |
| GBR | YRI | VB2 | ISAF (Equal -Ancestry) | 20% | 1.08 | 0.04 | 0.008 |
| GBR | YRI | VB2 | ISAF (Unequal-Ancestry) | 1% | 1.07 | 0.09 | 0.014 |
| GBR | YRI | VB2 | ISAF (Unequal-Ancestry) | 2% | 1.02 | 0.08 | 0.006 |
| GBR | YRI | VB2 | ISAF (Unequal-Ancestry) | 5% | 0.97 | 0.06 | 0.004 |
| GBR | YRI | VB2 | ISAF (Unequal-Ancestry) | 10% | 0.95 | 0.05 | 0.004 |
| GBR | YRI | VB2 | ISAF (Unequal-Ancestry) | 20% | 0.95 | 0.04 | 0.004 |
| CHS | GBR | VB1 | AFR | 1% | 0.07 | 0.04 | 0.868 |
| CHS | GBR | VB1 | AFR | 2% | 0.23 | 0.05 | 0.597 |
| CHS | GBR | VB1 | AFR | 5% | 0.40 | 0.04 | 0.366 |
| CHS | GBR | VB1 | AFR | 10% | 0.46 | 0.03 | 0.290 |
| CHS | GBR | VB1 | AFR | 20% | 0.47 | 0.02 | 0.282 |
| CHS | GBR | VB1 | EUR | 1% | 0.21 | 0.05 | 0.625 |
| CHS | GBR | VB1 | EUR | 2% | 0.40 | 0.05 | 0.364 |
| CHS | GBR | VB1 | EUR | 5% | 0.57 | 0.05 | 0.184 |
| CHS | GBR | VB1 | EUR | 10% | 0.66 | 0.04 | 0.119 |
| CHS | GBR | VB1 | EUR | 20% | 0.71 | 0.04 | 0.087 |
| CHS | GBR | VB1 | EAS | 1% | 1.24 | 0.11 | 0.069 |
| CHS | GBR | VB1 | EAS | 2% | 1.26 | 0.10 | 0.075 |
| CHS | GBR | VB1 | EAS | 5% | 1.27 | 0.08 | 0.077 |
| CHS | GBR | VB1 | EAS | 10% | 1.25 | 0.06 | 0.066 |
| CHS | GBR | VB1 | EAS | 20% | 1.18 | 0.05 | 0.035 |
| CHS | GBR | VB1 | Pooled | 1% | 0.36 | 0.06 | 0.409 |
| CHS | GBR | VB1 | Pooled | 2% | 0.55 | 0.06 | 0.207 |
| CHS | GBR | VB1 | Pooled | 5% | 0.71 | 0.05 | 0.086 |
| CHS | GBR | VB1 | Pooled | 10% | 0.79 | 0.05 | 0.047 |
| CHS | GBR | VB1 | Pooled | 20% | 0.83 | 0.04 | 0.031 |
| CHS | GBR | VB2 | ISAF (Equal-Ancestry) | 1% | 1.22 | 0.11 | 0.060 |
| CHS | GBR | VB2 | ISAF (Equal-Ancestry) | 2% | 1.23 | 0.10 | 0.060 |
| CHS | GBR | VB2 | ISAF (Equal -Ancestry) | 5% | 1.23 | 0.08 | 0.057 |
| CHS | GBR | VB2 | ISAF (Equal -Ancestry) | 10% | 1.20 | 0.06 | 0.044 |
| CHS | GBR | VB2 | ISAF (Equal -Ancestry) | 20% | 1.15 | 0.05 | 0.025 |
| CHS | GBR | VB2 | ISAF (Unequal-Ancestry) | 1% | 0.91 | 0.06 | 0.011 |
| CHS | GBR | VB2 | ISAF (Unequal-Ancestry) | 2% | 0.92 | 0.07 | 0.010 |
| CHS | GBR | VB2 | ISAF (Unequal-Ancestry) | 5% | 0.97 | 0.07 | 0.005 |
| CHS | GBR | VB2 | ISAF (Unequal-Ancestry) | 10% | 0.99 | 0.06 | 0.004 |
| CHS | GBR | VB2 | ISAF (Unequal-Ancestry) | 20% | 0.99 | 0.05 | 0.003 |
| CHS | CHS | VB1 | AFR | 1% | 0.02 | 0.03 | 0.956 |
| CHS | CHS | VB1 | AFR | 2% | 0.12 | 0.05 | 0.782 |
| CHS | CHS | VB1 | AFR | 5% | 0.25 | 0.03 | 0.562 |
| CHS | CHS | VB1 | AFR | 10% | 0.29 | 0.02 | 0.500 |
| CHS | CHS | VB1 | AFR | 20% | 0.29 | 0.01 | 0.500 |
| CHS | CHS | VB1 | EUR | 1% | 0.10 | 0.07 | 0.815 |
| CHS | CHS | VB1 | EUR | 2% | 0.24 | 0.05 | 0.573 |
| CHS | CHS | VB1 | EUR | 5% | 0.38 | 0.03 | 0.385 |
| CHS | CHS | VB1 | EUR | 10% | 0.42 | 0.02 | 0.338 |
| CHS | CHS | VB1 | EUR | 20% | 0.42 | 0.02 | 0.337 |
| CHS | CHS | VB1 | EAS | 1% | 0.90 | 0.10 | 0.020 |
| CHS | CHS | VB1 | EAS | 2% | 0.92 | 0.07 | 0.011 |
| CHS | CHS | VB1 | EAS | 5% | 0.95 | 0.06 | 0.006 |
| CHS | CHS | VB1 | EAS | 10% | 0.95 | 0.04 | 0.004 |
| CHS | CHS | VB1 | EAS | 20% | 0.94 | 0.03 | 0.004 |
| CHS | CHS | VB1 | Pooled | 1% | 0.19 | 0.08 | 0.659 |
| CHS | CHS | VB1 | Pooled | 2% | 0.34 | 0.05 | 0.435 |
| CHS | CHS | VB1 | Pooled | 5% | 0.48 | 0.04 | 0.275 |
| CHS | CHS | VB1 | Pooled | 10% | 0.52 | 0.03 | 0.233 |
| CHS | CHS | VB1 | Pooled | 20% | 0.53 | 0.02 | 0.222 |
| CHS | CHS | VB2 | ISAF (Equal-Ancestry) | 1% | 0.90 | 0.11 | 0.021 |
| CHS | CHS | VB2 | ISAF (Equal-Ancestry) | 2% | 0.91 | 0.07 | 0.012 |
| CHS | CHS | VB2 | ISAF (Equal -Ancestry) | 5% | 0.95 | 0.06 | 0.006 |
| CHS | CHS | VB2 | ISAF (Equal -Ancestry) | 10% | 0.95 | 0.04 | 0.004 |
| CHS | CHS | VB2 | ISAF (Equal -Ancestry) | 20% | 0.95 | 0.03 | 0.003 |
| CHS | CHS | VB2 | ISAF (Unequal-Ancestry) | 1% | 0.89 | 0.12 | 0.024 |
| CHS | CHS | VB2 | ISAF (Unequal-Ancestry) | 2% | 0.92 | 0.08 | 0.012 |
| CHS | CHS | VB2 | ISAF (Unequal-Ancestry) | 5% | 0.95 | 0.07 | 0.006 |
| CHS | CHS | VB2 | ISAF (Unequal-Ancestry) | 10% | 0.95 | 0.05 | 0.004 |
| CHS | CHS | VB2 | ISAF (Unequal-Ancestry) | 20% | 0.96 | 0.04 | 0.003 |
| CHS | YRI | VB1 | AFR | 1% | 0.09 | 0.09 | 0.828 |
| CHS | YRI | VB1 | AFR | 2% | 0.27 | 0.08 | 0.543 |
| CHS | YRI | VB1 | AFR | 5% | 0.45 | 0.05 | 0.302 |
| CHS | YRI | VB1 | AFR | 10% | 0.54 | 0.04 | 0.210 |
| CHS | YRI | VB1 | AFR | 20% | 0.61 | 0.03 | 0.155 |
| CHS | YRI | VB1 | EUR | 1% | 0.31 | 0.11 | 0.492 |
| CHS | YRI | VB1 | EUR | 2% | 0.51 | 0.08 | 0.250 |
| CHS | YRI | VB1 | EUR | 5% | 0.67 | 0.05 | 0.114 |
| CHS | YRI | VB1 | EUR | 10% | 0.72 | 0.04 | 0.083 |
| CHS | YRI | VB1 | EUR | 20% | 0.71 | 0.03 | 0.084 |
| CHS | YRI | VB1 | EAS | 1% | 1.38 | 0.14 | 0.161 |
| CHS | YRI | VB1 | EAS | 2% | 1.39 | 0.10 | 0.163 |
| CHS | YRI | VB1 | EAS | 5% | 1.38 | 0.07 | 0.151 |
| CHS | YRI | VB1 | EAS | 10% | 1.35 | 0.06 | 0.123 |
| CHS | YRI | VB1 | EAS | 20% | 1.24 | 0.04 | 0.061 |
| CHS | YRI | VB1 | Pooled | 1% | 0.47 | 0.13 | 0.298 |
| CHS | YRI | VB1 | Pooled | 2% | 0.66 | 0.09 | 0.123 |
| CHS | YRI | VB1 | Pooled | 5% | 0.82 | 0.06 | 0.036 |
| CHS | YRI | VB1 | Pooled | 10% | 0.89 | 0.05 | 0.015 |
| CHS | YRI | VB1 | Pooled | 20% | 0.93 | 0.04 | 0.007 |
| CHS | YRI | VB2 | ISAF (Equal-Ancestry) | 1% | 1.37 | 0.14 | 0.157 |
| CHS | YRI | VB2 | ISAF (Equal-Ancestry) | 2% | 1.38 | 0.11 | 0.154 |
| CHS | YRI | VB2 | ISAF (Equal -Ancestry) | 5% | 1.34 | 0.07 | 0.122 |
| CHS | YRI | VB2 | ISAF (Equal -Ancestry) | 10% | 1.29 | 0.06 | 0.085 |
| CHS | YRI | VB2 | ISAF (Equal -Ancestry) | 20% | 1.22 | 0.05 | 0.048 |
| CHS | YRI | VB2 | ISAF (Unequal-Ancestry) | 1% | 0.88 | 0.14 | 0.033 |
| CHS | YRI | VB2 | ISAF (Unequal-Ancestry) | 2% | 0.93 | 0.10 | 0.014 |
| CHS | YRI | VB2 | ISAF (Unequal-Ancestry) | 5% | 0.96 | 0.06 | 0.005 |
| CHS | YRI | VB2 | ISAF (Unequal-Ancestry) | 10% | 0.98 | 0.05 | 0.002 |
| CHS | YRI | VB2 | ISAF (Unequal-Ancestry) | 20% | 0.98 | 0.04 | 0.002 |
| YRI | GBR | VB1 | AFR | 1% | 1.43 | 0.23 | 0.236 |
| YRI | GBR | VB1 | AFR | 2% | 1.31 | 0.14 | 0.113 |
| YRI | GBR | VB1 | AFR | 5% | 1.28 | 0.08 | 0.081 |
| YRI | GBR | VB1 | AFR | 10% | 1.25 | 0.07 | 0.065 |
| YRI | GBR | VB1 | AFR | 20% | 1.17 | 0.06 | 0.034 |
| YRI | GBR | VB1 | EUR | 1% | 0.36 | 0.22 | 0.453 |
| YRI | GBR | VB1 | EUR | 2% | 0.46 | 0.19 | 0.329 |
| YRI | GBR | VB1 | EUR | 5% | 0.56 | 0.11 | 0.201 |
| YRI | GBR | VB1 | EUR | 10% | 0.62 | 0.07 | 0.153 |
| YRI | GBR | VB1 | EUR | 20% | 0.65 | 0.05 | 0.124 |
| YRI | GBR | VB1 | EAS | 1% | 0.32 | 0.20 | 0.504 |
| YRI | GBR | VB1 | EAS | 2% | 0.41 | 0.18 | 0.380 |
| YRI | GBR | VB1 | EAS | 5% | 0.52 | 0.11 | 0.245 |
| YRI | GBR | VB1 | EAS | 10% | 0.55 | 0.07 | 0.204 |
| YRI | GBR | VB1 | EAS | 20% | 0.56 | 0.04 | 0.198 |
| YRI | GBR | VB1 | Pooled | 1% | 0.57 | 0.26 | 0.248 |
| YRI | GBR | VB1 | Pooled | 2% | 0.67 | 0.17 | 0.138 |
| YRI | GBR | VB1 | Pooled | 5% | 0.76 | 0.10 | 0.066 |
| YRI | GBR | VB1 | Pooled | 10% | 0.80 | 0.07 | 0.043 |
| YRI | GBR | VB1 | Pooled | 20% | 0.83 | 0.05 | 0.030 |
| YRI | GBR | VB2 | ISAF (Equal-Ancestry) | 1% | 1.30 | 0.15 | 0.107 |
| YRI | GBR | VB2 | ISAF (Equal-Ancestry) | 2% | 1.25 | 0.11 | 0.072 |
| YRI | GBR | VB2 | ISAF (Equal -Ancestry) | 5% | 1.20 | 0.09 | 0.048 |
| YRI | GBR | VB2 | ISAF (Equal -Ancestry) | 10% | 1.15 | 0.07 | 0.028 |
| YRI | GBR | VB2 | ISAF (Equal -Ancestry) | 20% | 1.10 | 0.07 | 0.013 |
| YRI | GBR | VB2 | ISAF (Unequal-Ancestry) | 1% | 0.95 | 0.15 | 0.021 |
| YRI | GBR | VB2 | ISAF (Unequal-Ancestry) | 2% | 0.93 | 0.08 | 0.012 |
| YRI | GBR | VB2 | ISAF (Unequal-Ancestry) | 5% | 0.93 | 0.08 | 0.011 |
| YRI | GBR | VB2 | ISAF (Unequal-Ancestry) | 10% | 0.93 | 0.06 | 0.008 |
| YRI | GBR | VB2 | ISAF (Unequal-Ancestry) | 20% | 0.94 | 0.06 | 0.006 |
| YRI | CHS | VB1 | AFR | 1% | 1.32 | 0.13 | 0.120 |
| YRI | CHS | VB1 | AFR | 2% | 1.30 | 0.11 | 0.098 |
| YRI | CHS | VB1 | AFR | 5% | 1.26 | 0.07 | 0.071 |
| YRI | CHS | VB1 | AFR | 10% | 1.22 | 0.05 | 0.049 |
| YRI | CHS | VB1 | AFR | 20% | 1.14 | 0.04 | 0.022 |
| YRI | CHS | VB1 | EUR | 1% | 0.35 | 0.20 | 0.455 |
| YRI | CHS | VB1 | EUR | 2% | 0.46 | 0.18 | 0.319 |
| YRI | CHS | VB1 | EUR | 5% | 0.56 | 0.10 | 0.204 |
| YRI | CHS | VB1 | EUR | 10% | 0.59 | 0.06 | 0.168 |
| YRI | CHS | VB1 | EUR | 20% | 0.60 | 0.03 | 0.159 |
| YRI | CHS | VB1 | EAS | 1% | 0.29 | 0.17 | 0.528 |
| YRI | CHS | VB1 | EAS | 2% | 0.40 | 0.18 | 0.395 |
| YRI | CHS | VB1 | EAS | 5% | 0.51 | 0.11 | 0.252 |
| YRI | CHS | VB1 | EAS | 10% | 0.56 | 0.06 | 0.194 |
| YRI | CHS | VB1 | EAS | 20% | 0.61 | 0.03 | 0.157 |
| YRI | CHS | VB1 | Pooled | 1% | 0.55 | 0.24 | 0.259 |
| YRI | CHS | VB1 | Pooled | 2% | 0.66 | 0.16 | 0.136 |
| YRI | CHS | VB1 | Pooled | 5% | 0.75 | 0.09 | 0.068 |
| YRI | CHS | VB1 | Pooled | 10% | 0.79 | 0.04 | 0.044 |
| YRI | CHS | VB1 | Pooled | 20% | 0.83 | 0.02 | 0.030 |
| YRI | CHS | VB2 | ISAF (Equal-Ancestry) | 1% | 1.29 | 0.13 | 0.098 |
| YRI | CHS | VB2 | ISAF (Equal-Ancestry) | 2% | 1.25 | 0.11 | 0.074 |
| YRI | CHS | VB2 | ISAF (Equal -Ancestry) | 5% | 1.20 | 0.07 | 0.043 |
| YRI | CHS | VB2 | ISAF (Equal -Ancestry) | 10% | 1.15 | 0.04 | 0.023 |
| YRI | CHS | VB2 | ISAF (Equal -Ancestry) | 20% | 1.10 | 0.04 | 0.011 |
| YRI | CHS | VB2 | ISAF (Unequal-Ancestry) | 1% | 0.94 | 0.15 | 0.023 |
| YRI | CHS | VB2 | ISAF (Unequal-Ancestry) | 2% | 0.95 | 0.09 | 0.010 |
| YRI | CHS | VB2 | ISAF (Unequal-Ancestry) | 5% | 0.93 | 0.05 | 0.007 |
| YRI | CHS | VB2 | ISAF (Unequal-Ancestry) | 10% | 0.93 | 0.03 | 0.005 |
| YRI | CHS | VB2 | ISAF (Unequal-Ancestry) | 20% | 0.94 | 0.03 | 0.004 |
| YRI | YRI | VB1 | AFR | 1% | 0.95 | 0.09 | 0.010 |
| YRI | YRI | VB1 | AFR | 2% | 0.92 | 0.06 | 0.009 |
| YRI | YRI | VB1 | AFR | 5% | 0.89 | 0.05 | 0.014 |
| YRI | YRI | VB1 | AFR | 10% | 0.89 | 0.04 | 0.013 |
| YRI | YRI | VB1 | AFR | 20% | 0.89 | 0.03 | 0.012 |
| YRI | YRI | VB1 | EUR | 1% | 0.22 | 0.13 | 0.619 |
| YRI | YRI | VB1 | EUR | 2% | 0.29 | 0.14 | 0.518 |
| YRI | YRI | VB1 | EUR | 5% | 0.36 | 0.09 | 0.423 |
| YRI | YRI | VB1 | EUR | 10% | 0.38 | 0.06 | 0.391 |
| YRI | YRI | VB1 | EUR | 20% | 0.36 | 0.03 | 0.405 |
| YRI | YRI | VB1 | EAS | 1% | 0.18 | 0.11 | 0.680 |
| YRI | YRI | VB1 | EAS | 2% | 0.25 | 0.13 | 0.575 |
| YRI | YRI | VB1 | EAS | 5% | 0.32 | 0.09 | 0.474 |
| YRI | YRI | VB1 | EAS | 10% | 0.34 | 0.06 | 0.438 |
| YRI | YRI | VB1 | EAS | 20% | 0.33 | 0.03 | 0.452 |
| YRI | YRI | VB1 | Pooled | 1% | 0.36 | 0.18 | 0.433 |
| YRI | YRI | VB1 | Pooled | 2% | 0.44 | 0.14 | 0.333 |
| YRI | YRI | VB1 | Pooled | 5% | 0.49 | 0.08 | 0.267 |
| YRI | YRI | VB1 | Pooled | 10% | 0.51 | 0.05 | 0.242 |
| YRI | YRI | VB1 | Pooled | 20% | 0.51 | 0.03 | 0.245 |
| YRI | YRI | VB2 | ISAF (Equal-Ancestry) | 1% | 0.94 | 0.10 | 0.012 |
| YRI | YRI | VB2 | ISAF (Equal-Ancestry) | 2% | 0.92 | 0.06 | 0.011 |
| YRI | YRI | VB2 | ISAF (Equal -Ancestry) | 5% | 0.88 | 0.04 | 0.016 |
| YRI | YRI | VB2 | ISAF (Equal -Ancestry) | 10% | 0.88 | 0.04 | 0.015 |
| YRI | YRI | VB2 | ISAF (Equal -Ancestry) | 20% | 0.88 | 0.03 | 0.015 |
| YRI | YRI | VB2 | ISAF (Unequal-Ancestry) | 1% | 0.94 | 0.08 | 0.010 |
| YRI | YRI | VB2 | ISAF (Unequal-Ancestry) | 2% | 0.92 | 0.07 | 0.011 |
| YRI | YRI | VB2 | ISAF (Unequal-Ancestry) | 5% | 0.88 | 0.04 | 0.016 |
| YRI | YRI | VB2 | ISAF (Unequal-Ancestry) | 10% | 0.88 | 0.04 | 0.015 |
| YRI | YRI | VB2 | ISAF (Unequal-Ancestry) | 20% | 0.88 | 0.03 | 0.015 |

